# Transcriptomic characterization of signaling pathways associated with osteoblastic differentiation of MC-3T3E1 cells

**DOI:** 10.1101/410597

**Authors:** Louis M. Luttrell, Moahad S. Dar, Diane Gesty-Palmer, Hesham M. El-Shewy, Katherine M. Robinson, Courtney J. Haycraft, Jeremy L. Barth

**Affiliations:** Department of Medicine, Medical University of South Carolina, Charleston, SC, USA; Department of Biochemistry & Molecular Biology, Medical University of South Carolina, Charleston, SC, USA; Ralph H. Johnson Veterans Affairs Medical Center, Charleston, SC, USA; Department of Medicine, Duke University Medical Center, Durham, NC, USA; Department of Biology, Mississippi College, Clinton, MS, USA; Department of Regenerative Medicine and Cell Biology, Medical University of South Carolina, Charleston, SC, USA

## Abstract

Bone remodeling involves the coordinated actions of osteoclasts, which resorb the calcified bony matrix, and osteoblasts, which refill erosion pits created by osteoclasts to restore skeletal integrity and adapt to changes in mechanical load. Osteoblasts are derived from pluripotent mesenchymal stem cell precursors, which undergo differentiation under the influence of a host of local and environmental cues. To characterize the autocrine/paracrine signaling networks associated with osteoblast maturation and function, we performed gene network analysis using complementary “agnostic” DNA microarray and “targeted” NanoString^™^ nCounter datasets derived from murine MC3T3-E1 cells induced to undergo synchronized osteoblastic differentiation *in vitro*. Pairwise datasets representing changes in gene expression associated with growth arrest (day 2 to 5 in culture), differentiation (day 5 to 10 in culture), and osteoblast maturation (day 10 to 28 in culture) were analyzed using Ingenuity Systems^™^ Pathways Analysis to generate predictions about signaling pathway activity based on the temporal sequence of changes in target gene expression. Our data indicate that some pathways known to be involved in osteoblast differentiation, e.g. Wnt/β-catenin signaling, are most active early in the process, while others, e.g. TGFβ/BMP, cytokine/JAK-STAT and TNFα/RANKL signaling, increase in activity as differentiation progresses. Collectively, these pathways contribute to the sequential expression of genes involved in the synthesis and mineralization of extracellular matrix. These results provide insight into the temporal coordination and complex interplay between signaling networks controlling gene expression during osteoblast differentiation. A more complete understanding of these processes may aid the discovery of novel methods to promote osteoblast development for the trea™ent of conditions characterized by low bone mineral density.

## INTRODUCTION

Bone remodeling is the continuous process through which worn bone is removed and replaced [1, 2]. Bone-resorbing osteoclasts differentiate from hematopoietic stem cell precursors in response to cues originating from osteocytes, bone lining cells, and differentiating osteoblasts. Bone-forming osteoblasts derive from mesenchymal stem cell presursors and undergo a defined maturational sequence from proliferating preosteoblasts to mature synthetically active osteoblasts, before finally undergoing apoptosis or transforming into osteocytes embedded within the bony matrix and quiescent bone lining cells covering the mineralized surface. The bone remodeling cycle involves sequential osteoclastic bone resorption followed by the synthesis and mineralization of new bone matrix by osteoblasts, a process that requires several weeks to complete. Since these osteoclast-osteoblast bone forming units that mediate this process are constantly being created and destroyed, any analysis performed on bone tissue, whether by classical histomorphometry or using genomic and proteomic methods, is a ‘snapshot’ of the metabolic state of bone at that moment in time. While such *in vivo* studies are extremely useful for understanding the effects of disease, hormone administration/withdrawal or drug trea™ent on overall bone metabolism, they inevitably capture cross sectional data from multiple cell types in different differentiation states.

In contrast, *in vitro* studies offer the advantage that cellular development can be synchronized, offering a better opportunity to view differentiation as a linear process. In bone, the replication of undifferentiated osteogenic precursor cells, their recrui™ent to remodeling bone matrix, and their subsequent acquisition of differentiated function, results from the complex interplay of signals transmitted by mechanical load, polypeptide growth factors, steroid and thyroid hormones, and locally produced cytokines and prostaglandins [3, 4]. While circulating hormones play an important modulatory role, osteoblastic differentiation can be induced *in vitro*, indicating that, once triggered, the process is autonomous, i.e. independent of ongoing exposure to systemically derived factors.

Gene array technology is a potentially powerful tool for understanding complex biological processes. A significant limitation of the approach, however, is that it is difficult to translate lists of significantly regulated genes into changes in biologically relevant signaling networks. Genomic datasets are invariably incomplete and contain some number of false positive ‘hits’, making candidate based follow up studies unreliable. In addition, important pathway components may not be regulated at the transcriptional level. Circumventing these limitations requires the use of bioinformatic approaches that compare changes in gene expression against databases of known protein-protein interactions to establish the probability that a given signaling or metabolic pathway is regulated under varying experimental conditions [5, 6]. These *in silico* analyses, which enable gene expression profile data to be expressed as the statistical probability that a particular pathway is regulated, can “fill in the blanks”, leading to a more holistic view of process-related changes in signaling pathway activity.

To better understand the temporal regulation of osteoblast differentiation, we performed microarray analysis of gene expression followed by signal transduction pathways analysis on murine MC3T3-E1 cells undergoing osteoblastic differentiation *in vitro*. Taking advantage of their well-defined maturational sequence [7-10], we isolated RNA at four stages: during log growth, early and late osteoblastic differentiation, and mature synthetic function. We then performed pairwise comparisons to identify significant changes in gene expression associated with each of these stages of osteoblast development, and used the resulting genesets to identify the time-dependent changes in signal transduction pathway activity. Our data indicate that the temporally coordinated activation of signaling pathways known to be involved in osteoblast differentiation, e.g. Wnt/β-catenin, Transforming Growth Factor-β (TGFβ) Bone Morphogenic Protein (BMP), cytokine/Janus Kinase (JAK)-Signal Transducer and Activator of Transcription (STAT), and Tumor Necrosis factor-α (TNFα/Receptor Activator of Nuclear Factor κ-B (NFκB) Ligand (RANKL) signaling, correlates with the sequential expression of genes involved in the biosynthesis and mineralization of extracellular matrix as differentiation progresses. These results demonstrate the utility of functional genomic approaches to microarray analysis and offer insight into the temporal sequence of changes in the autocrine/paracrine signaling networks regulating osteoblast differentiation.

## Materials and methods

### Culture and differentiation of MC3T3-E1 cells

Stock cultures of MC3T3-E1 cells (subclone 4; CRL-2593; ATCC) were maintained in α-minimum essential medium (MEM) supplemented with 10% v/v fetal bovine serum, penicillin (100 units/mL) and streptomycin (100 pg/mL) in a humidified 10% CO_2_ a™osphere at 37°C. Until the time of study cells were maintained in log phase growth by passage every 3-5 days using 0.001% pronase (w/v) to detach adherent cells. For studies of the temporal sequence of osteoblast differentiation, cells were plated at an initial density of 20,000 cells/well in 6-well plates or 100,000 cells/dish in 10 cm dishes, and grown for 2 to 28 days in α-MEM supplemented with 10% v/v fetal bovine serum, 5 mM β-glycerol phosphate and 50 µg/mL ascorbic acid (7-10).

### Cell replication

Between days 1 and 5 in culture, cells in 6-well plates were treated with 0.001% pronase (w/v) to achieve detachment and directly counted in a hemocytometer.

### Alkaline phosphatase activity

Alkaline phosphatase activity was measured by para-nitrophenyl phosphate hydrolysis as previously described (11). Briefly, MC3T3-E1 cells growth in 6-well plates were harvested in distilled water and disrupted by sonication. Appropriately diluted aliquots of cell lysate containing equal cell protein were incubated for 30 min at 37 °C in a final reaction volume of 600 L, containing 1.0 M diethanolamine, pH 10.3 and 15 mM para-nitrophenyl phosphate. Reactions were terminated by the addition of 2.4 mL 0.1N NaOH, after which generation of para-nitrophenol was measured by determining absorbance at 400 nm. Results were expressed as pmol para-nitrophenol/min/10^6^ cells.

### Synthesis of type I collagen

Type 1 collagen production was determined by western blotting. Monolayers of MC3T3-E1 cells were lysed directly in 1X Laemmli sample buffer, dispersed by sonication, and resolved by sodium dodecyl sulfate – polyacrylamide gel electrophoresis. Immune complexes on nitrocellulose membranes were detected using mouse monoclonal anti-type I collagen IgG_1_ (COL1A: sc59772; Santa Cruz Biotechnology, Santa Cruz, CA) with horseradish peroxidase-conjugated donkey anti-mouse IgG (Code: 715-035-150; Jackson ImmunoResearch Laboratories Inc., West Grove, PA) as secondary antibody. The cell content of each sample was determined by western blotting in parallel for α-actin using mouse monoclonal anti-actin IgG_1_ (C-2: sc8432; Santa Cruz Biotechnology, Santa Cruz, CA). Immune complexes were visualized on X-ray film by enzyme-linked chemiluminescence and quantified using a Fluor-S MultiImager. Data were expressed as the ratio of type I collagen to α-actin in each sample.

### Alizarin red staining

Matrix mineralization was quantified by Alizarin red staining as described (12). Monolayers of MC3T3-E1 cells in 6-well plates were fixed for 24 hr in a 10% formalin:methanol:distilled water solution (1:1:1.5), stained for 20 min in 2% Alizarin Red-S in distilled water, washed with distilled water and air dried. Mineralization was quantified by eluting the stain using 10% cetylpyridium chloride and measuring absorbance at 520 nM.

### mRNA isolation

MC3T3-E1 cells were cultured as described in 10 cm dishes for 2, 5, 10 or 28 days prior to isolation of RNA. Total RNA from three independent cultures was isolated at each time point. Cells were harvested by scraping and RNA was isolated with Trizol Reagent (Invitrogen, Carlsbad, CA) and purified using the RNeasy kit (Qiagen Inc., Valencia, CA) according to the manufacturer’s protocols (13). Total RNA was analyzed for concentration (ng/L) and purity (ratios of 260/280 nm and 260/230 nm) using a NanoDrop 1000 Spectrophotometer (Thermo Scientific, Wilmington, DE). RNA integrity was analyzed using the Experion RNA HighSens Analysis Kit (Bio-Rad Laboratories Inc., Hercules CA).

### Microarray analysis

Samples underwent RNA amplification (Message Amp; Ambion, Austin, TX), labeling with Cy3, and hybridization to Mouse Operon 17,000 gene feature (Operon dataset; version 2.0) spotted oligonucleotide arrays in the microarray facility of the Duke University Institute for Genome Sciences and Policy (*www.genome.duke.edu/cores/microarray/*). MIAME compliant microarray data files have been deposited with the NCBI GEO database (*www.ncbi.nlm.nih.gov/gds*) (GEO Series GSE64485). Data pre-processing and normalization were performed on GenePix scan results files (.gpr files) using the Bioconductor LimmaGUI package 1.28.0 run with R 2.13.0 software (14). Background correction was performed using the normexp method with offset of 16, and spot quality weighting was applied as follows: 1 for Good (100) or Unflagged (0); 0.1 for Bad (−100), Not Found (−50) and Absent (−75) flags. Print-tip group loess normalization was applied for normalization within arrays. Review of box plots of normalized M values indicated that normalization between arrays was not warranted. Normalized M values, i.e. log2 test(Cy3)/reference(Cy5) relationship, were imported into dchip for comparative analysis. ANOVA was used to find genes differing as function of time, i.e. significantly different between any two time points. ANOVA filtering at the 0.005 level yielded 1005 genes passing with a reasonable 5-10% false discovery rate (17664 compared; expected false positive: 88). Self-organizing maps (SOM) were used to partition the significantly regulated genes into different response patterns. Expression data were imported into MeV software for SOM analysis, z-standardization performed, i.e. mean=0 and SD=1, and SOM clusters for ANOVA p<0.005 were generated by: 16 clusters, 4×4, 2000 iterations, hexagonal topography, Gaussian neighborhood, alpha 0.05, radius 1.0, no HCL linkage, Pearson correlation.

### NanoString^™^ nCounter analysis

The NanoString^™^ nCounter gene expression system (NanoString^™^ Technologies; Seattle, WA) was used for expression profiling of selected mRNA species isolated from MC3T3-E1 cells at Days 2, 5, 10 and 28 in culture using a custom nCounter CodeSet composed of 243 probes (S1 Table) including 6 housekeeping controls (Eif4a2, GusB, Oaz1, Stk36, Tceb1, and Tubb4a). With NanoString^(tm)^ technology fluorescent single strand RNA probes are hybridized to complimentary target strands of mRNA and quantified based on the fluorescence of each target gene within each sample (15,16). Briefly, the NanoString^™^ reporter probe CodeSet was suspended in 70µL of hybridization buffer and 8µL aliquots were combined in sterile microfuge tubes with each RNA sample diluted to a concentration of 250ng RNA in 5µL. Thereafter, 2µL of the capture probe CodeSet was added to each tube, tubes were centrifuged, and then incubated at 65°C in a BioRad T100 Thermal Cycler (BioRad; Hercules, CA) for 13-15 hours. After hybridization, samples were analyzed using the NanoString^™^ Technologies Prep Station and Digitial Analyzer according to manufacturer’s instructions. All 12 samples, i.e. RNA from triplicate cultures at each of four time points, were analyzed simultaneously to minimize batch effects. The resulting counts were analyzed using NanoStriDE and GraphPad Prism 7 (GraphPad Software; Carlsbad, CA) software. Statistical significance of change over time was determined by two-way ANOVA with Tukey’s multiple comparisons test using GraphPad Prism 7.

### IPA metabolic pathways analysis

Network analysis of genesets representing changes in mRNA abuundance between specified time points was performed using the Ingenuity Systems^™^ Pathways Analysis (IPA) tool (Qiagen; Redwood City, CA). IPA compares Genbank Accession number/expression information with a proprietary protein-interaction database to establish the probability that a given signaling or metabolic pathway is activated under varying experimental conditions. For the microarray dataset, expression ratios for all relevant pairwise comparisions, i.e. D2 vs D5, D2 vs D10, D2 vs D28, D5 vs D10, D5 vs D28, and D10 vs D28, were calculated using the ANOVA p<0.005 set of 1005 significantly regulated genes. For the NanoString^™^ dataset, pairwise expression ratios were calculated for each of the 237 measurable genes. Expression ratio data were uploaded into the IPA Pathways Analysis system (*https://analysis.ingenuity.com/*), yielding 976 analyzable transcripts from the microarray dataset (S2 Table). Each dataset was subjected to IPA Core Analysis, then analyzed using IPA Upstream Regulator, Downstream Effects, and Canonical Pathways analytic tools. To capture pathway changes associated with each phase of differentiation we focused on the D2 vs D5, D5 vs D10, and D10 vs D28 pairwise comparisons. For IPA Upstream Regulator and Canonical Pathways analysis, gene clusters composed of ≥ 2 genes per group with *P* < 0.05 enrichment, i.e. –log(p-value) ≥ 1.3, compared with a standard murine background database were considered analyzable. The IPA output was exported as Microsoft Excel files to prepare the S3-7 Tables. Graphic representations of the data were prepared using either the IPA Canonical Pathways Molecular Activity Predictor tool or GraphPad Prism 7 software, as appropriate. To facilitate visual inspection of the changes in predicted Upstream Regulator and Canonical Pathways activity associated with each interval, heat maps were generated from the activation z-scores using Morpheus software (*https://software.broadinstitute.org/morpheus/*).

## Results

### The temporal sequence of MC3T3-E1 cell differentiation

MC3T3-E1 cells undergo a well-characterized process of osteoblastic differentiation when placed in culture medium supplemented with β-glycerol phosphate and ascorbic acid [7-10]. Fig 1 presents this process tracked using traditional markers: cell number, bone alkaline phosphatase, abundance of type I collagen, and alizarin red staining. After initial seeding, the cells remain in log phase growth for 2-3 days, undergoing growth arrest upon attaining confluence by days 3-4. Osteoblastic differentiation begins upon growth arrest and continues through days 5 to 10 in culture, evident first as an increase in the production of bone-specific alkaline phosphatase, followed by deposition of a collagenous matrix composed in part of type 1 collagen. Matrix mineralization begins as early as day 10 and accelerates with time in culture. By day 28 the MC3T3-E1 derived osteoblasts have produced a mineralized matrix.

**Fig 1.**
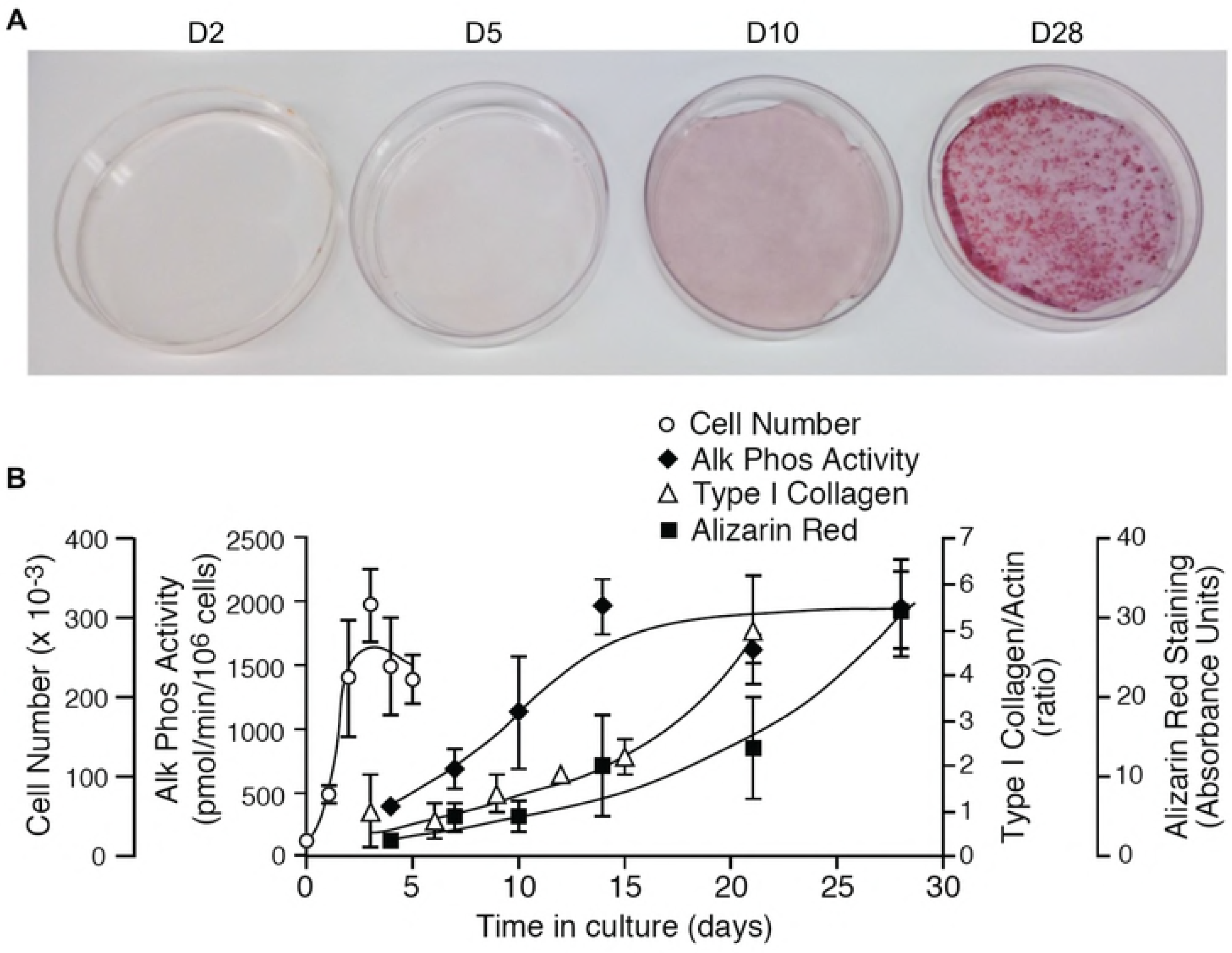
MC3T3-E1 osteoblast maturation *in vitro*. MC3T3-E1 cells were seeded in 6-well tissue culture plates at an initial density of 20,000 cells/well and maintained in culture for up to 28 days. **A.** Representative Alizarin Red stained culture dishes from Days 2, 5, 10 and 28 demonstrating the progression of matrix mineralization. **B.** Graph depicting change in cell number (days 1-5), secreted alkaline phosphatase activity (days 4-28), type 1 collagen synthesis (days 3-21), and matrix mineralization (days 4-28) associated with MC3T3-E1 differentiation. Data shown are the Mean ± SEM of triplicate determinations. These data were used to select time points representing proliferating preosteoblasts (day 2), early and late differentiating osteoblasts (days 5 and 10), and active osteoblasts (day 28), for subsequent mRNA isolation.

Predictably, osteoblastic differentiation on MC3T3-E1 cells is reflected in changes in the abundance of mRNA encoding bone marker proteins. As shown in Fig 2, osteoblast developmental markers, matrix components, and proteins involved in cell adhesion and matrix remodeling change over time as the cells evolve from proliferating pre-osteoblasts to mature osteoblasts. Notably, these changes in mRNA abundance appear at different times during development. mRNA encoding Runx2, the first transcription factor required for determination of the osteoblast lineage [17,18], increases early in development and plateaus between Days 5 and 10, while others, e.g. alkaline phosphatase (*Alp1*), integrin-binding sialoprotein (*Ibsp*), a major structural protein of the bone matrix, and parathyroid hormone receptor (*Pthr1*), increase steadily from Day 2 to Day 28. Still other mRNA species are abundant throughout development, e.g. collagen type 1A (*Col1a1*), and some increase between Days 10 and 28 after differentiation is well underway e.g. the osteoblast-specific matrix protein periostin (*Postn*). Such differences are consistent with a temporally coordinated process wherein early events trigger the sequential activation of a transcriptional program driven by intracellular signaling networks.

**Fig 2.**
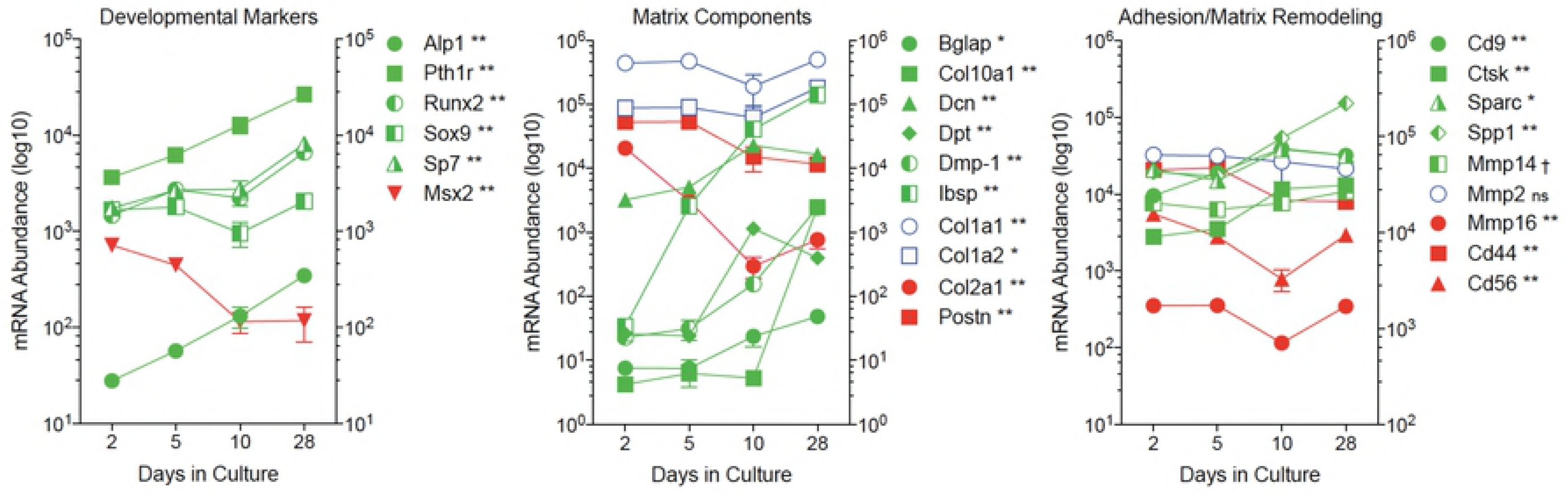
Temporal changes in the abundance of mRNA encoding bone marker proteins. Total RNA was isolated from triplicate cultures of MC3T3-E1 cells at Days 2, 5, 10 and 28 in culture, and mRNA abundance quantified by NanoString nCounter using a bone focused probe set (S1 Table). Developmental markers shown are: alkaline phosphatase (*Alp1*); parathyroid hormone receptor (*Pthr1*); the transcription factors *Runx2, Sox9* and *Sp7*; and the transcriptional repressor *Msx2*. Matrix components shown are: bone gamma-carboxyglutamate protein (*Bglap*); collagen types 1A1 (*Col1a1*), 1A2 (*Col1a2*), 2A1 (*Col2a1*) and 10A1 (*Col10a1*); decorin (*Dcn*); dermatopontin (*Dpt*); dentin matrix protein-1 (*Dmp-1*); integrin-binding sialoprotein (*Ibsp*); and periostin (*Postn*). Proteins associated with cell adhesion and matrix remodeling are: tetraspanin (*Cd9*); cathepsin K (*Ctsk*); osteonectin (*Sparc*); osteopontin (*Spp1*); matrix metalloproteinases 2 (*Mmp2*), 14 (*Mmp14*), and 16 (*Mmp16*); hyaluronic acid receptor (*Cd44*); and neural cell adhesion molecule 1 (*Cd56*). Data shown represent the Mean ± SD of triplicate samples. Error bars not shown are smaller than the symbol. † P < 0.05; ^*^ P < 0.01; ^**^ P < 0.001 different in abundance between at least two time points by two-way ANOVA with Tukey’s multiple comparisons test; ns, not significant.

### DNA microarray analysis of MC3T3-E1 differentiation

DNA microarrays, because they capture information about the abundance of a large number of unselected mRNA species, provide an “agnostic” snapshot of gene expression patterns at a given point in time. Combining microarray data on changes in mRNA abundance over time with bioinformatic tools, such as Ingenuity Systems^™^ IPA, provides a means to translate microarray data into a more complete picture of metabolic activity [5, 6]. To identify changes in gene expression occurring at different stages of differentiation, triplicate samples of total mRNA were isolated from subconfluent MC3T3-E1 preosteoblasts (day 2), growth-arrested preosteoblasts (day 5), differentiating osteoblasts (day 10) and maturing synthetically-active osteoblasts (day 28), and hybridized to Operon V2.0 murine cDNA microarrays representing approximately 17,600 expressed sequence tags. Raw microarray data (GEO Series GSE64485) were analyzed by ANOVA to identify genes whose mean expression was significantly different between any two time points. Fig 3A shows a heat map of 1005 mRNAs passing the ANOVA filtered at p < 0.005. S2 Table lists the gene symbol, annotation, and observed abundance of the 976 analyzable mRNAs from this dataset. Hierarchical clustering revealed several distinct temporal patterns of expression, with some gene clusters increasing or decreasing in abundance early in differentiation, others changing progressively throughout differentiation, or changing most dramatically during the period of osteoblast maturation. Still others genes exhibited a biphasic pattern, increasing or decreasing with the onset of differentiation and reversing their direction of change between days 10 and 28 in culture. To further partition genes into different response patterns, we generated self-organizing maps (SOM) from the ANOVA p<0.005 dataset. As shown in Fig 3B, distinct temporal patterns of mRNA abundance were evident, reflecting each stage of osteoblast differentiation.

**Fig 3.**
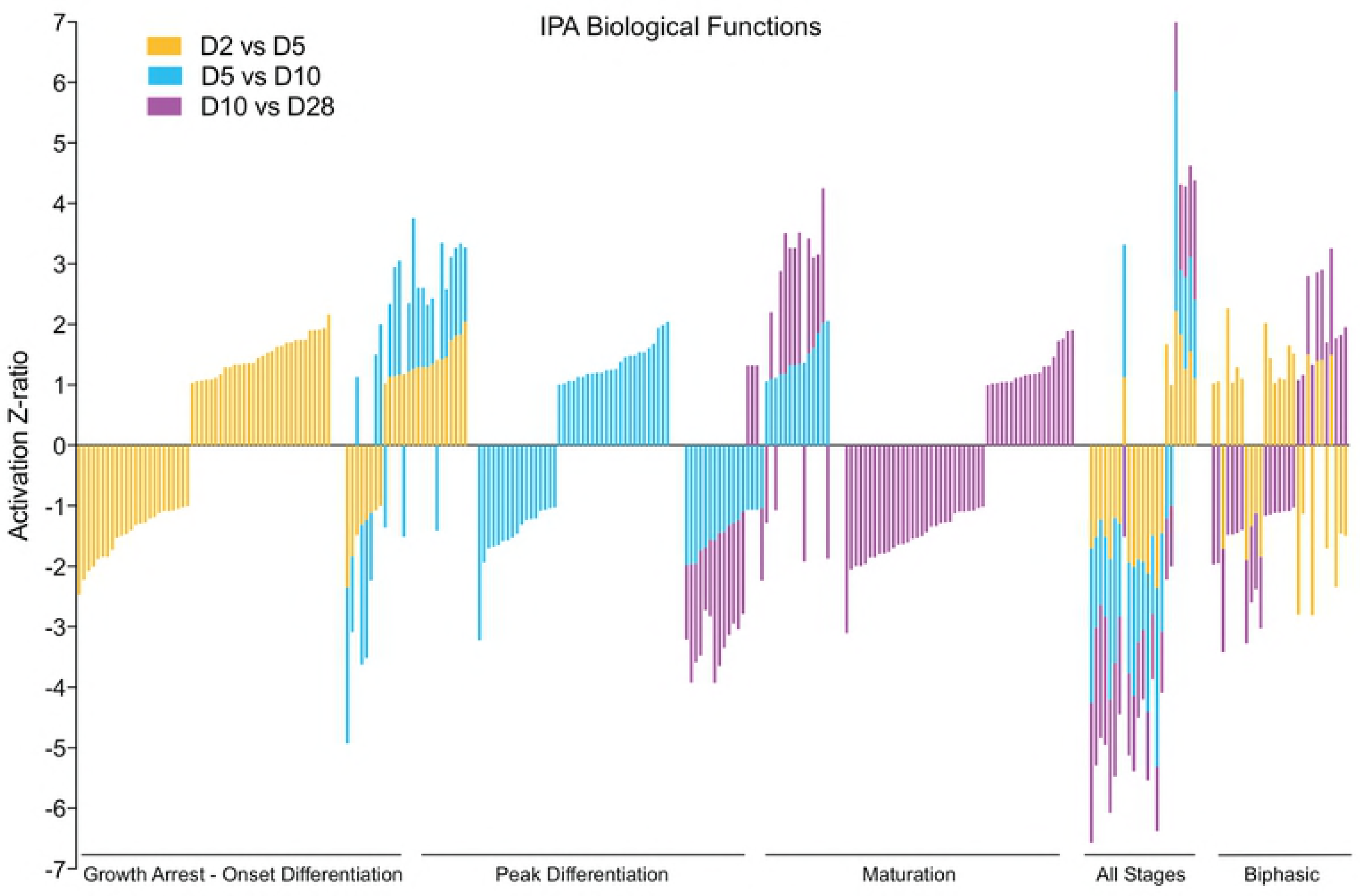
Temporal patterns of change in the MC3T3-E1 transcriptome during differentiation. Triplicate DNA microarrays at each time point were used to identify significantly regulated mRNAs at different phases of osteoblast differentiation by ANOVA (p<0.005; estimated false discovery rate 8.8%). **A.** Heat map representing observed mRNA abundance of 1005 genes identified by ANOVA as demonstrating a significant difference between any two time points. Hierarchical clustering was used to identify coordinated patterns of change. **B.** Sixteen cluster SOM representing temporal changes in mRNA abundance associated with MC3T3-E1 differentiation. Expression data were subjected to z-standardization and SOM assembled using MeV software. The resulting 16 SOM clusters are shown grouped in relation to the differentiation state of MC3T3-E1 cells. Growth arrest was associated with abrupt changes (increase or decrease) in mRNA levels between days 2 and 5 (240 genes). The onset of differentiation was associated with progressive changes in mRNA levels between days 2 and 10 (212 genes). Peak differentiation was associated with prominent changes in mRNA levels between days 5 and 10 (246 genes). Osteoblast maturation was associated with prominent changes in mRNA levels between days 10 and 28 (307 genes).

To test the hypothesis that the biological processes underlying osteoblastic differentiation of MC3T3-E1 cells are reflected in the coordinated changes in the transcriptome over time, the DNA microarray data were analyzed using Ingenuity Systems^™^ IPA software. IPA compares empirically derived “omics” datasets, e.g. DNA microarray data, with a curated database of reported gene-gene and protein-protein interactions to predict signaling pathway activity based on observed changes in upstream regulators and/or the downstream genes whose expression they control. The IPA output includes two statistical measures. The first, which is typically expressed as –log(p-value), represents the probability that the correlation between an input set of observed factors and co-regulated genesets in the IPA database did not occur by chance. Hence, a –log(p-value) greater than 1.3 represents p < 0.05 of a significant association. The second, termed an activation z-score, is based on the degree to which observed changes in factor levels, e.g. increases or decreases in mRNA abundance between two points in time, correlate with the expected changes associated with pathway activation or inhibition. Thus, an activation z-score > 2 or < -2 predicts pathway activation or inhibition, respectively, with p < 0.05.

The biochemical characterization of differentiating MC3T3-E1 cells (Fig 1) demonstrates that the major downstream biological processes, e.g. cell proliferation versus matrix mineralization, change over time. To generate a gestalt view of whether the structure of the DNA microarray dataset reflects this temporal evolution, we calculated expression ratios for each of the 976 analyzable genes identified by ANOVA using three pairwise comparisons, day 2 to day 5 (D2 vs D5), day 5 to day 10 (D5 vs D10), and day 10 to day 28 (D10 vs D28), and performed IPA Downstream Effects Analysis, which predicts increases or decreases in downstream biological activities. S3 Table lists the annotation, -log(p-value), and activation z-score for all biological process terms identified from our dataset where the z-score was > 1 or < -1. These results are presented graphically in Fig 4. As shown, each pairwise comparison was associated with a set of unique of terms, here represented graphically as vertical bars. Importantly, terms identified in two overlapping comparisons exhibited a high degree of concordance in the predicted direction of activation/inhibition (20 of 26 terms appearing in both the D2 vs D5 and D5 vs D10 comparison, and 24 of 31 terms appearing in both the D5 vs D10 and the D10 vs D28 comparison). Consistent with the SOM analysis (Fig 3B), where some gene clusters increased in abundance steadily throughout differentiation, several process level terms were identified in all three genesets, and again there was strong concordance in the predicted direction of activation/inhibition (20 of 23 terms appearing in all three comparisons). Of interest, process terms appearing only in the D2 vs D5 and D10 vs D28 comparisons showed less concordance (only 9 of 29 terms were concordant). This too may reflect at the process level expression patterns observed in the SOM analysis, where some gene clusters clearly underwent reciprocal regulation, increasing/decreasing between days 2 and 5, remaining relatively constant between days 5 and 10, and returning to their prior levels between days 10 and 28.

**Fig 4.**
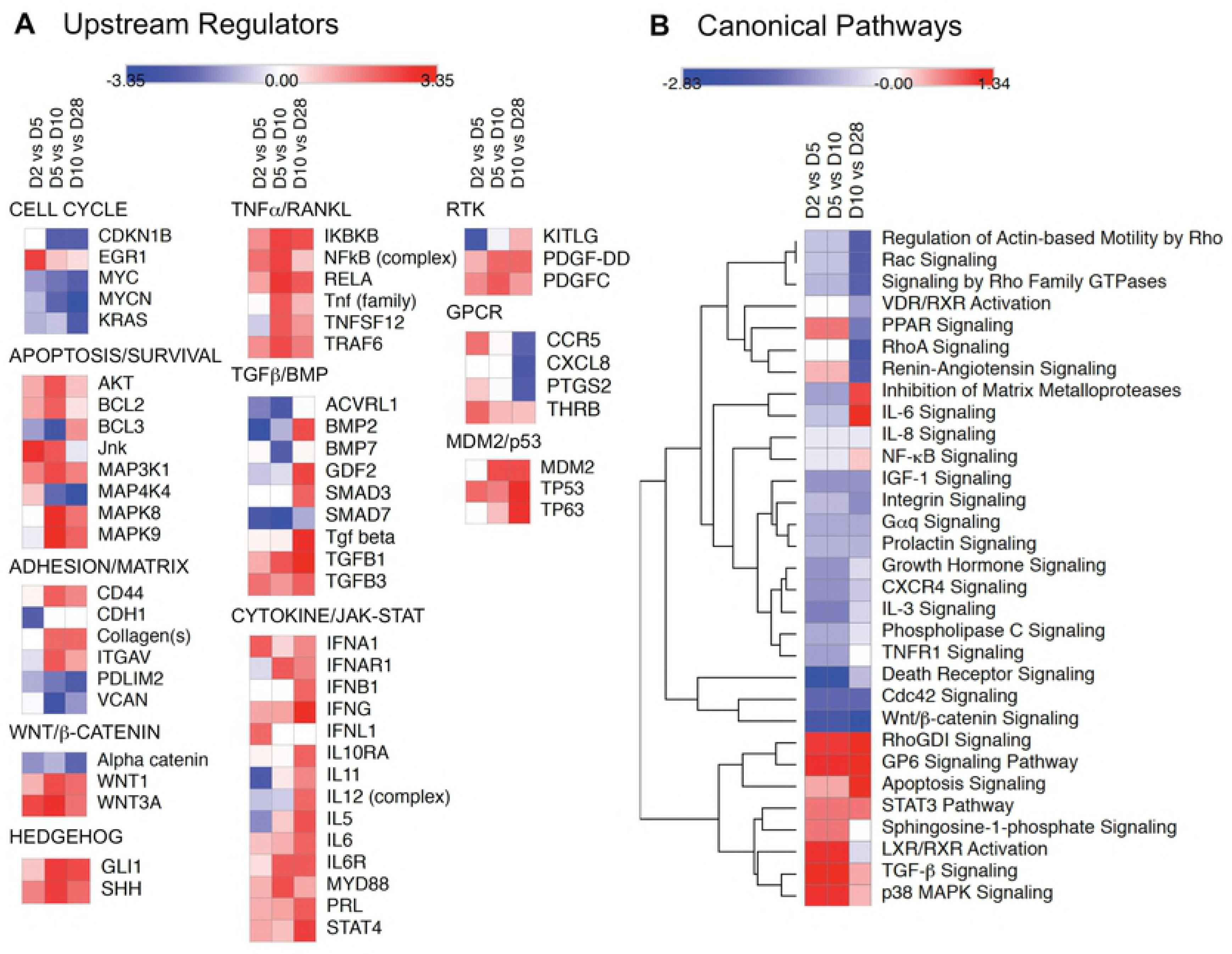
Temporal changes in mRNA abundance reflect evolving biological processes during MC3T3-E1 differentiation. The mRNA abundance of 976 significantly regulated genes identified by ANOVA as changing during MC3T3-E1 differentiation was used to calculate expression ratios comparing D2 vs D5, D5 vs D10, and D10 vs D28. For each pairwise comparison, the earlier time point was used as the denominator and later time point as the numerator, such that expression ratios reflect increases/decreases in mRNA abundance as differentiation proceeds. IPA Downstream Effects Analysis was performed to identify biological process terms associated with each interval and filtered to include terms only with –log(p value) >1.3, minimum of two genes, and z-score >1 or <-1. The graph depicts z-score values for terms associated with the period of growth arrest and onset of differentiation (gold bars), active differentiation (blue bars), and osteoblast maturation (lavender bars). The descriptive annotations associated with each term are omitted for simplicity but presented in S3 Table.

To resolve the temporal changes in signaling networks associated with osteoblastic differentiation of MC3T3-E1 cells we performed IPA Upstream Regulator and Canonical Pathways Analysis using the 976 significantly regulated genes identified by ANOVA. The IPA Upstream Regulator Analysis predicts which transcriptional regulators are activated or inhibited based on observed changes in expression of downstream genes. Predicted upstream regulators with activation Z-scores > 2 or < -2 during at least one phase of differentiation are shown in S4 Table. Individual upstream regulators were grouped based on the signaling networks with which they are most associated, and the z-scores derived from the D2 vs D5, D5 vs D10, and D10 vs D28 comparisons used to generate heat maps that illustrate the predicted change in regulator activity as differentiation progresses. In these maps, rows represent individual upstream regulators and columns represent time intervals. Predicted increases in activity from the beginning to end of each interval, e.g. from day 2 to day 5, are indicated in red and decreases in blue, with color intensity representing the magnitude of the z-score. Thus, an upstream regulator that was predicted to increase in activity from day 2 to day 5, day 5 to day 10, and day 10 to day 28 would be red in all columns, while one that increased from day 2 to day 5 and then remained active at the same level would be red in the D2 vs D5 column, then white in the D5 vs D10 and D10 vs D28 columns. As shown in Fig 5A, upstream regulators associated with cell cycle progression were predicted to become less active over time, consistent with the growth arrest of MC3T3-E1 cells that heralds the onset of differentiation. Conversely, upstream regulators of several pathways associated osteoblast differentiation, e.g. TGFβ/BMP/SMAD, WNT/β-catenin, and Hedgehog signaling [4] were predicted to become more active as differentiation progressed, as did regulators of TNFα/RANKL/NFκB and cytokine/JAK-STAT signaling.

**Fig 5.**
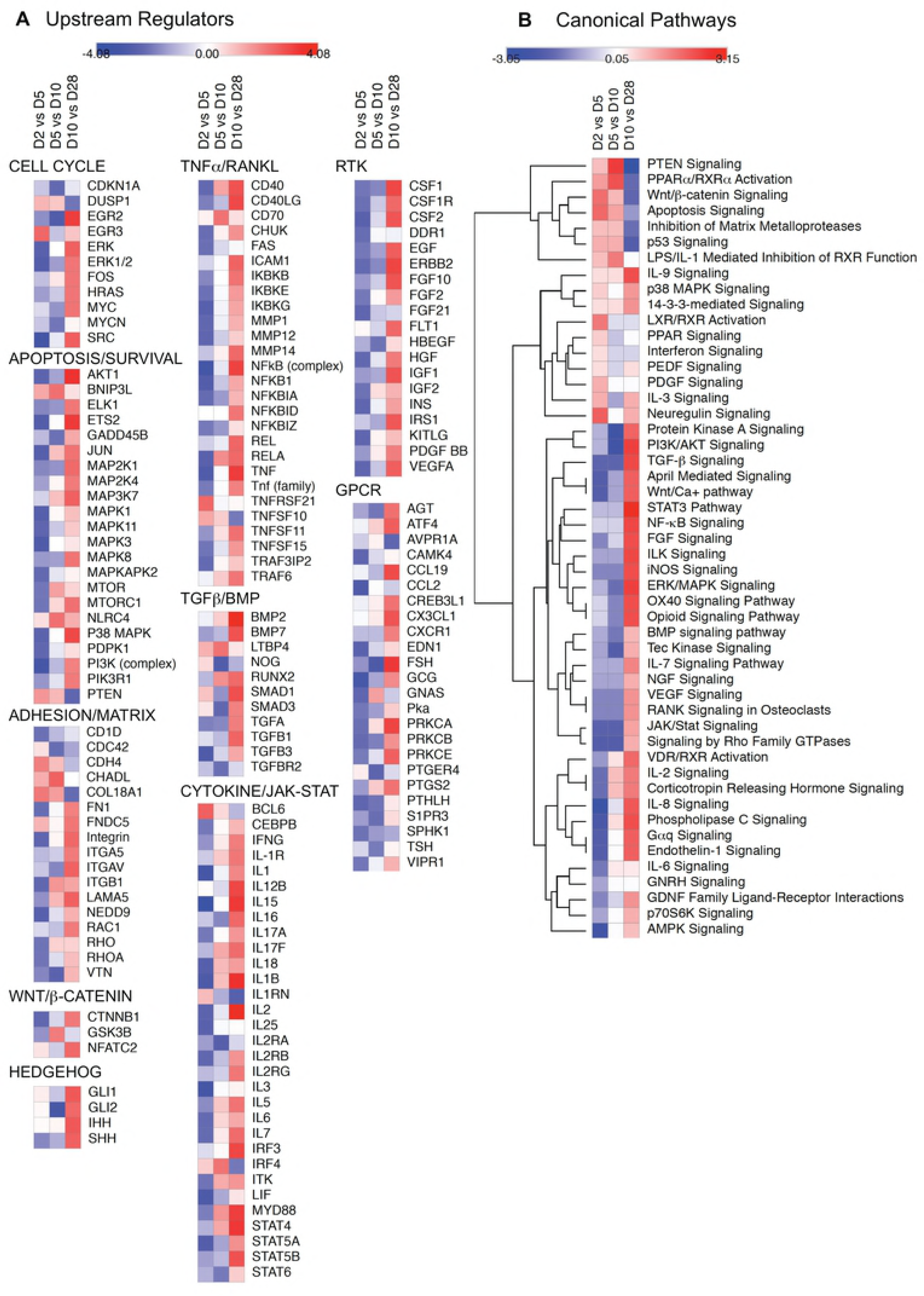
Temporal changes in predicted upstream regulators and canonical signaling pathways associated with MC3T3-E1 cell differentiation. Microarray data on the 976 significantly regulated mRNA species were used to calculate change in expression ratio between D2 vs D5, D5 vs D10, and D10 vs D28. Expression ratios were analyzed using IPA Upstream Regulator and Canonical Pathways Analysis software and heat maps reflecting the changes in predicted activity during each interval were generated using Morpheus software. **A.** Heat maps depicting changes in selected upstream regulators (rows) with activation z-scores > 2 (red) or < -2 (blue) during at least one phase of differentiation (columns). Upstream regulators were arbitrarily grouped based on their involvement is biological processes or signaling pathways related to osteoblast differentiation. **B.** Heat maps depicting changes in z-score for selected canonical signaling pathways (rows) during each phase of differentiation (columns). Z-scores were subjected to Euclidean hierarchical clustering in Morpheus to group pathways based on similarity in temporal change.

To gain insight into how predicted changes in the activity of individual upstream regulators were integrated into signaling networks, we next performed IPA Canonical Pathways Analysis, which compares observed changes in mRNA abundance to the expected direction of change associated with pathway activation or inhibition. The list of signaling pathways represented in the dataset along with –log(p-value) and activation z-score are shown in S5 Table. Fig 5B depicts a heat map of predicted changes in activity in selected canonical signaling pathways. Hierarchical clustering was performed to group pathways based on similarities in the change in activity over time. Considering pathways known to be involved in bone development, Wnt/β-catenin signaling decreased in activity as differentiation progressed, while TGFβ signaling increased. The TGF-β/BMP axis is a principal regulator of mesenchymal stem cell differentiation into cartilage and bone [19-22], acting through several effectors including SMADs, p38 mitogen-activated protein kinase (MAPK), and phosphatidyl inositol 3-kinase (PI3K)/AKT. TGF-β/BMP engages in extensive cross talk with other receptor-mediated signaling in bone, including WNT/β- catenin, Notch, Hedgehog, fibroblast growth factor (FGF), parathyroid hormone-related peptide (PTHrp), and interleukin (IL)/TNFα/interferon-γ cytokines that collectively signal via the JAK/STAT and NFκB pathways [4,23]. Notably, several of these pathways, e.g. p38 MAPK, STAT3, NFκB and IL6 signaling also showed a trend toward activation during differentiation.

### NanoString^™^ analysis of MC3T3-E1 differentiation

To validate our “agnostic” microarray data on signaling pathway activation, we performed a “focused” analysis of MC3T3-E1 cell gene rexpression using NanoString^™^ nCounter. The NanoString^™^ nCounter system uses color-coded molecular “barcodes” attached target-specific probes to count up to several hundred unique transcripts in a single hybridization reaction [15,16]. The culture protocol used for the microarray experiment was repeated to provide independent mRNA samples. Triplicate samples of total mRNA isolated from MC3T3-E1 cells at days 2, 5, 10 and 28 in culture were analyzed using a NanoString^™^ Code Set designed to quantify the abundance of 237 transcripts related to bone development and signaling. S1 Table lists the gene name, annotation, and expression data for the NanoString^™^ probes. Fig 6A shows a heat map of all 237 transcripts assayed. As with the microarray data, hierarchical clustering revealed several distinct temporal patterns of expression, with some groups of transcripts increasing/decreasing in abundance early in differentiation and others changing most dramatically later during osteoblast maturation. Fig 6B-E shows temporal changes in selected transcripts related to pathways identified in the bioinformatics analysis of the microarray data. Significant changes in mRNA abundance were detected in ligands, receptors or modulators of BMP, TGFβ and Activin signaling, the three closely-related components of the TGFβ network, as well as in the TNFα-NFκB, interleukin-JAK/STAT, and WNT/β-catenin pathways.

**Fig 6.**
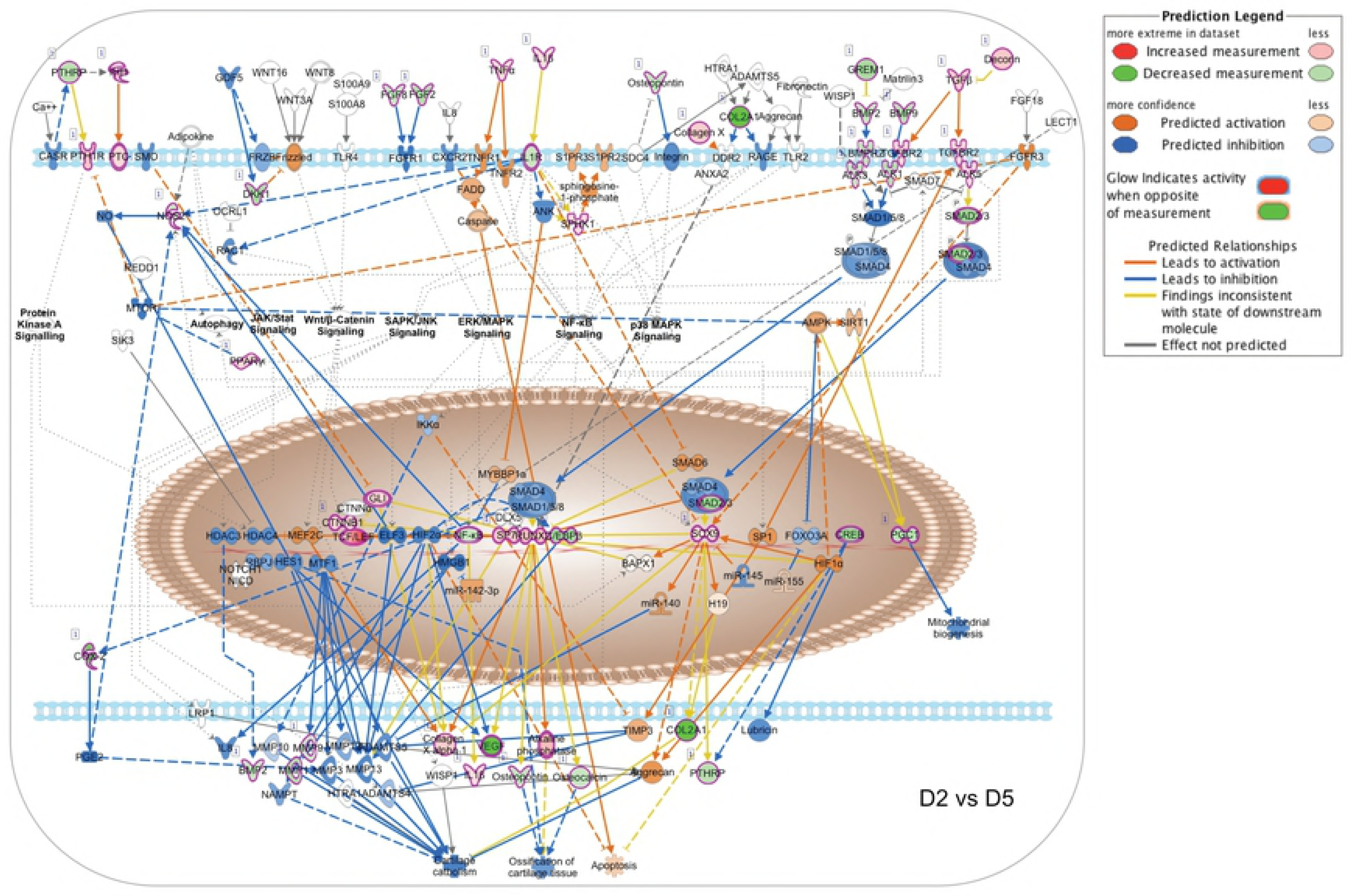
NanoString^™^ analysis of bone-related mRNAs during MC3T3-E1 cell differentiation. Total RNA was isolated from triplicate cultures of MC3T3-E1 cells at days 2, 5, 10 and 28, and mRNA abundance quantified by NanoString^™^ nCounter using a bone specific Code Set (S1 Table). **A.** Heat map depicting changes in mRNA abundance for individual mRNA species (rows) over time in culture (columns) for day 2 (D2), day 5 (D5), day 10 (D10), and day 28 (D28). Expression data, after log2 adjus™ent, were subjected to Euclidean heirarchical clustering in Morpheus to group genes based on similarity in temporal change. **B.** mRNA abundance of selected ligands, receptors, modulators, and mediators related to BMP/TGFβ/Activin, TNFα/NFκB, IL/JAK-STAT, and WNT/β-catenin signaling. BMP pathway components shown are: BMP 4 (*Bmp4*); BMP receptor 1A (*Bmpr1a*); BMP receptor 2 (*Bmpr2*); the BMP co-receptors, repulsive guidance molecule (RGM) A (*Rgma*) and RGM B (*Rgmb*); the BMP negative regulators, Chordin and Noggin; and the DAN family BMP antagonist, Gremlin. TGFβpathway components shown are: TGFβ1 (*Tgfb1*); TGFβ2 (*Tgfb2*); TGFβ3 (*Tgfb3*); TGFβ receptor 1 (*Tgfbr1*); and TGFβ receptor 2 (*Tgfbr2*). Activin pathway components shown are: inhibin subunit βA (*Inhba*); activin A receptor type 1 (*Acvr1*); activin A receptor type 1B (*Acvr1b*); activin A receptor type 2A (*Acvr2a*); BMP and activin membrane bound inhibitor (*Bambi*); and the activin and TGFβ receptor ligand, left-right determination factor 1 (*Lefty*). TNFα pathway components shown are: TNF ligand superfamily member 13-like (*April*); TNF (*Tnf*); RANKL (*Tnfsf11*); TNF-receptor superfamily member 4 (*Tnfrsf4*); receptor activator of NFκB (*Tnfrsf11a*); TNF receptor superfamily member 11b (*Tnfrsf11b*); and NFκB (*Nfkb*). Interleukin pathway components shown are: IL1B (*Il1b*); IL4 (*Il4*); IL7 (*Il7*); IL12A (*Il12a*); IL1 receptor-like 1 (*Il1rl1*); IL2 receptor β subunit (*Il2rb*); IL4 receptor α subunit (*Il4ra*); IL15 receptor α subunit (*Il15ra*); and STAT1 (*Stat1*). WNT pathway components shown are: WNT 5A (*Wnt5a*); Wnt 7A (*Wnt7a*); the WNT signaling pathway inhibitor, Dickkopf (*Dkk1*); β-catenin (*Ctnnb1*); the regulator of β-catenin stability, Axin 2 (*Axin2*); and the β-catenin regulated transcription factors, nuclear factor of activated T cells 1 (*Nfatc1*) and transcription factor 7 (*Tcf7*). In each graph, symbols representing ligands are show in green, receptor subunits in blue, intracellular mediators and modulators in red, and transcription factors in lavender. Data shown represent the Mean ± SD of triplicate samples. Error bars not shown are smaller than the symbol. † P<0.05; ^*^ P<0.01; ^**^ P<0.001 different in abundance between at least two time points by two-way ANOVA with Tukey’s multiple comparisons test; ns, not significant.

To determine how the changing levels of pathway components translated into changes in pathway activity during differentiation, we performed IPA Upstream Regulator and Canonical Pathways Analysis using expression ratios derived from comparisons of the NanoString^™^ data for D2 vs D5, D5 vs D10, and D10 vs D28. Predicted upstream regulators with activation z-scores > 2 or < -2 during at least one phase of differentiation are shown in S6 Table. Selected upstream regulators were grouped based on the signaling networks with which they are most associated, and the z-scores used to generate heat maps. As shown in Fig 7A, upstream regulator activity associated with cell cycle progression, apoptosis, and cell survival tended to decrease between days 2 and 5 and days 5 and 10, then increase between days 10 and 28. Notably, activity of the anti-apoptotic regulators AKT1 and p38 MAPK that function downstream of TGFβ/BMP and TNFα/RANKL increased as differentiation progressed. Coincident with this, upstream regulators related to TGFβ/BMP/SMAD, WNT/β-catenin, and Hedgehog signaling showed activation during osteoblastic differentiation, as did regulators involved in TNFα/RANKL/NFκB, cytokine/JAK-STAT, receptor tyrosine kinase (RTK), and G protein-coupled receptor (GPCR) signaling.

**Fig 7.**
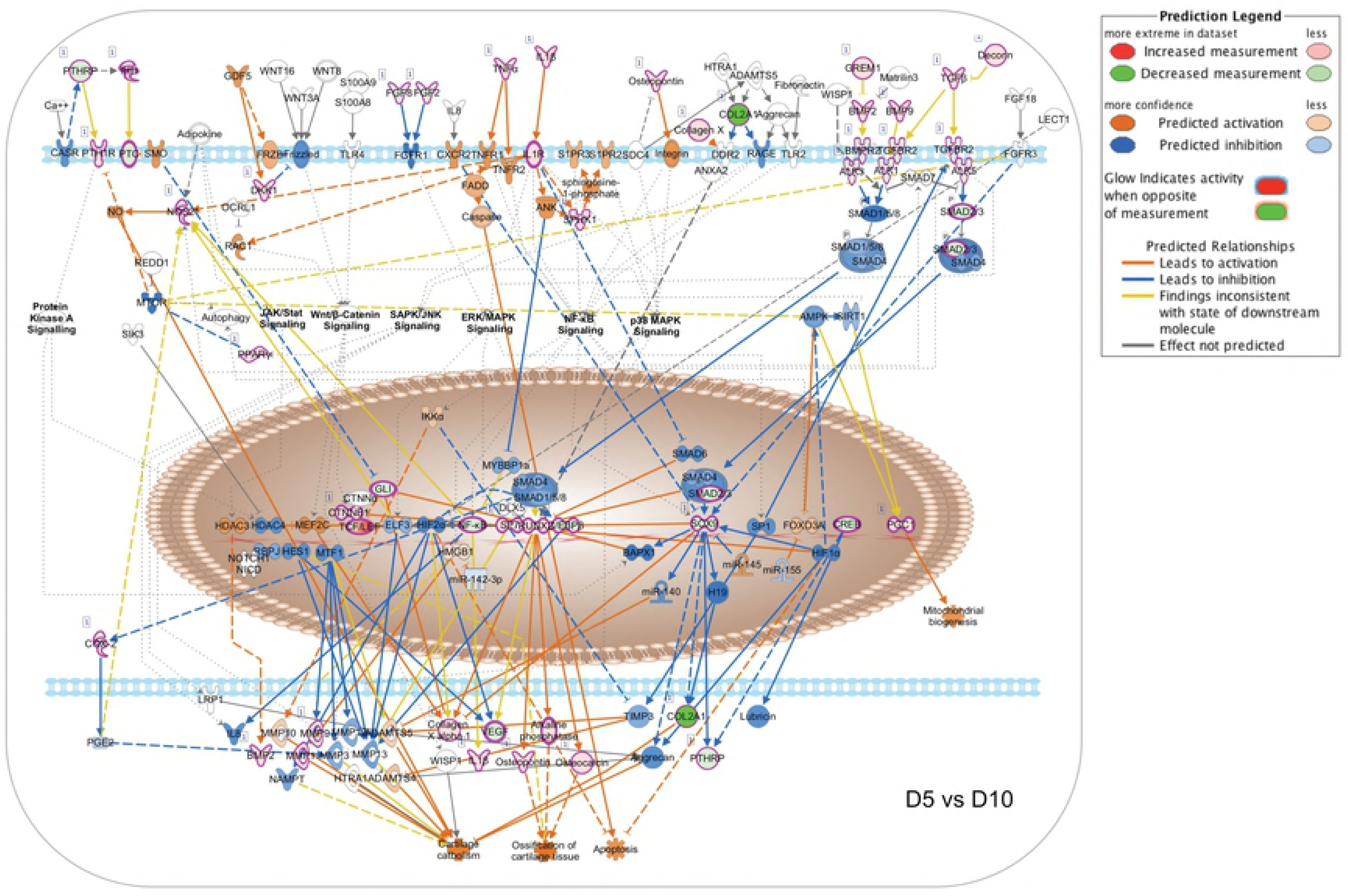
Upstream regulators and canonical signaling pathways analysis of a focused NanoString^™^ dataset. NanoString^™^ nCounter data on the abundance of 237 bone-related mRNA species were used to calculate change in expression ratio between D2 vs D5, D5 vs D10, and D10 vs D28. Expression ratios were analyzed using IPA Upstream Regulator and Canonical Pathways Analysis software and heat maps reflecting the changes in predicted activity during each interval were generated using Morpheus software. **A.** Heat maps depicting changes in selected upstream regulators (rows) with activation z-scores > 2 (red) or < -2 (blue) during at least one phase of differentiation (columns). Upstream regulators were arbitrarily grouped based on their relationship to biological processes or signaling pathways involved in osteoblast differentiation. **B.** Heat maps depicting changes in z-score for selected canonical signaling pathways (rows) during each phase of differentiation (columns). Z-scores were subjected to Euclidean hierarchical clustering in Morpheus to group pathways based on similarity in temporal change.

We next performed IPA Canonical Pathways Analysis using the NanoString^™^ dataset. The list of signaling pathways represented along with –log(p-value) and activation z-score are shown in S7 Table. Fig. 7B depicts a heat map of predicted changes in activity in selected canonical signaling pathways. Hierarchical clustering was performed to group pathways based on similarities in the change in activity over time. Consistent with the canonical pathways analysis of the microarray dataset (Fig 5), WNT/β-catenin signaling became less active as differentiation progressed, while TGFβ and BMP signaling increased, although the changes in the NanoString^™^ dataset were more apparent later than in the microarray dataset, between day 10 and day 28. Interleukin signaling was predicted to increase progressively between days 5 and 28, with JAK-STAT signaling showing the activation between days 10 and 28. TNFα/NFκB pathway signaling was likewise predicted to increase in both the NanoString^™^ and microarray datasets.

### Overview of signaling networks during MC3T3-E1 differentiation

To test the overall similarity between the NanoString^™^ and microarray datasets we compared the pathway activity predictions generated from each using the IPA Canonical Pathways molecular activity predictor tool, which graphically depicts the predicted change in pathway activity based on observed changes in upstream and downstream gene expression. S1 Fig shows the WNT/β-catenin pathway comparison using D2 vs D5 expression ratios from each dataset. Since WNT/β-catenin signaling was predicted to decline as differentiation progressed, it would be most active during this interval. Consistent with this, both datasets indicated β-catenin pathway activation in this time frame, as well as inhibition of the negative regulatory TGFβ/TGFβ-activated kinase 1 (TAK1)/p38 MAPK/nemo like kinase (NLK) input from the TGFβ receptor pathway that was predicted to be less active early in differentiation. S2 Fig compares IPA predicted changes in activity within the TGFβ/BMP signaling network occurring between days 10 and 28, an interval during which both datasets indicated pathway activation. While the focused Nanostring^™^ dataset better captured activation of BMP receptor signaling during this phase of differentiation, both datasets predicted net activation of the SMAD2/3 and TAK1/p38 MAPK components of TGFβ signaling. S3 Fig compares the IPA predicted changes in the canonical TNFα/NFκB signaling pathway between days 10 and 28 in culture. The TNFα network plays a key role during osteoblast maturation, acting as an inhibitor of osteoblast differentiation and, along with RANKL, promoting osteoclast development [24,25]. Both datasets indicated net activation of NFκB-dependent transcription during MC3T3-E1 maturation related to changes in the expression of TNF family and growth factor ligands and receptors. Both datasets also indicated relative inhibition of interleukin receptor-mediated NFκB activation through TNF receptor-associated factor 6 (TRAF6)/TAK1. S4 Fig illustrates the predicted activation of canonical STAT3 signaling downstream of cytokine and growth factor receptors between days 10 and 28 observed in both the microarray and Nanostring^™^ nCounter datasets. Collectively, the data indicate substantial concordance between the two independent MC3T3-E1 datasets and highlight the evolving changes in WNT/β- catenin, TGFβ/BMP/SMAD, TNFα/RANKL/NFκB, and cytokine/JAK-STAT signaling associated with osteoblast differentiation.

To illustrate the temporal evolution of signaling network interaction during MC3T3-E1 cell differentiation, we generated pathway activity predictions for the IPA osteoarthritis canonical pathway, which integrates multiple signal inputs controlling expression of bone-related genes. As the first transcription factor required for osteoblastic differentiation, control of Runx2-dependent transcription is central to the process [17,18]. Runx2 activity reflects the input of multiple upstream regulators, notably including BMP receptors signaling via SMAD1/5/8 as well as TGFβ and activin receptors signaling through SMAD2/3. Given that the NanoString^™^ Code Set was selected to examine bone-related genes, the osteoarthritis network was the most heavily populated canonical pathway in our IPA analysis with a -log(p-value) of 43.3 (S7 Table). Fig 8 shows the pathway activity analysis based on D2 vs D5 expression ratio changes from the NanoString^™^ nCounter dataset. During this phase, MC3T3-E1 cells are transitioning from log phase growth to growth arrest and initiating the process of differentiation. Based on observed upregulation of Runx2, Sp7 and Sox9 mRNA, Runx2-dependent transcription is predicted to increase from Day 2 to Day 5, associated with increases in mRNA encoding collagen species and alkaline phosphatase. Observed changes in genes encoding Indian Hedgehog (IHH), Patched (PTCH) and β-catenin, as well as TNFα, IL1B, TGFβ and TGFβ receptor 2 (TGFBR2), suggest that the onset of differentiation coincides with upregulation of autocrine ligands and receptors that later come to drive the differentiation process. Notably, TGFβ/BMP signaling is not yet predicted to be active due to relative downregulation of BMP2/9 and SMAD2/3.

**Fig 8.**
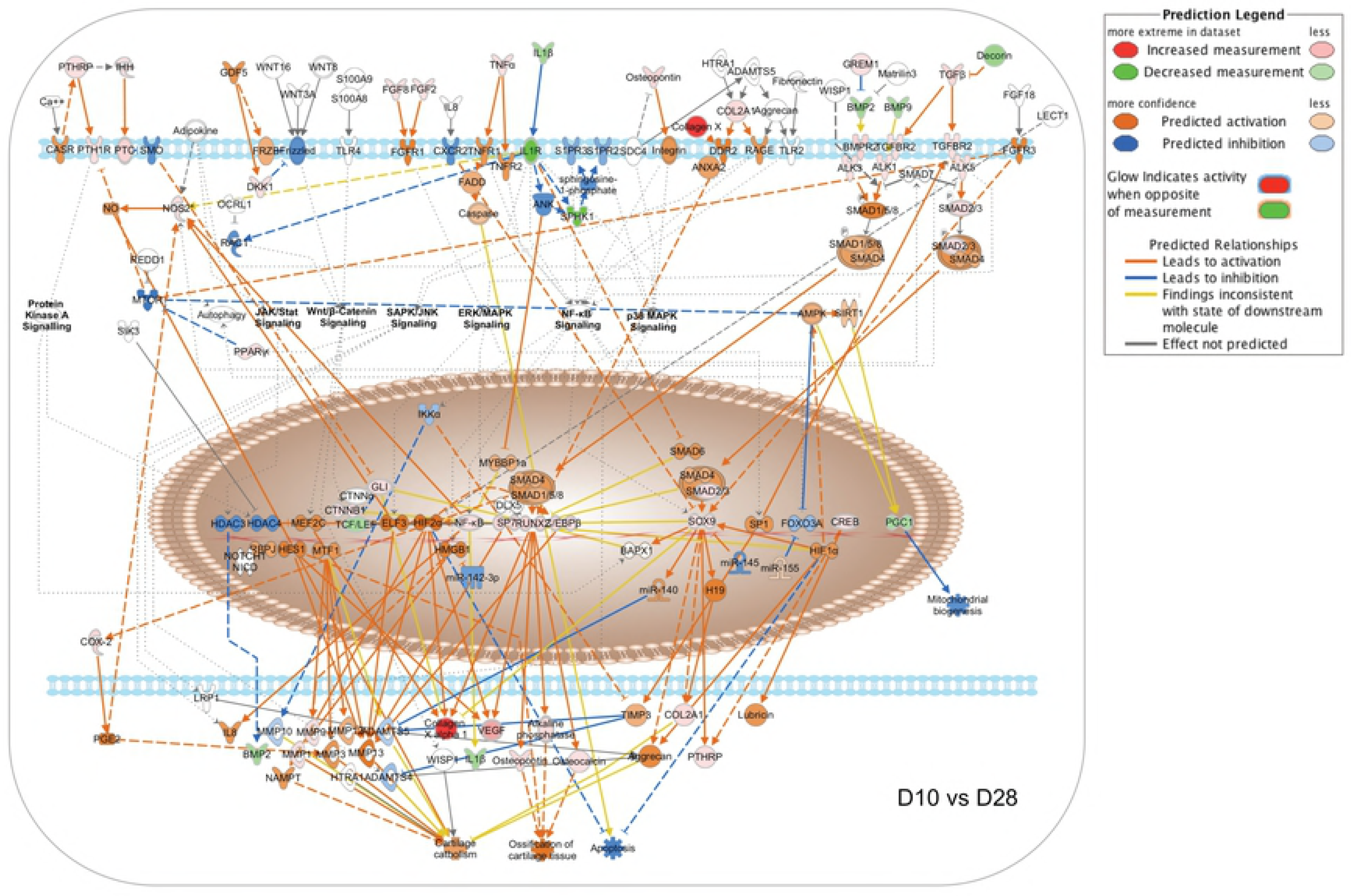
Changes in canonical signaling pathway activity in MC3T3-E1 cells between days 2 and 5. Expression ratios representing the changing abundance of 237 bone-related mRNA species in MC3T3-E1 cells between days 2 and 5 in culture were used to populate the IPA osteoarthritis pathway network and signaling pathway activation state was assessed using the IPA molecular pathway predictor tool. As indicated in the prediction legend, observed upregulation and downregulation of mRNAs are shown in red and green, respectively, while predicted activation or inhibition of signaling intermediates and pathways are shown in orange and blue.

Fig 9 depicts the results of an identical analysis performed using the D5 vs D10 expression ratio changes from the Nanostring^™^ nCounter dataset. This phase is associated with osteoblastic differentiation and increased expression of secreted growth factors and matrix components, but relatively little matrix mineralization (Figs 1 and 2). The network analysis suggests that increasing expression of Runx2 and Sp7 is now associated with upregulation of the Runx2-regulated matrix components osteopontin (Spp1) and osteocalcin (Bglap2), and further increases in expression of collagen species and alkaline phosphatase, along with the upstream regulators IHH, β-catenin, TNFα, IL1B, and TGFβ. Increasing expression of BMP2 and BMP9 is now evident, although the molecular pathway predictor still suggests that SMAD1/5/8 and SMAD2/3 signaling is attenuated. Fig 10 depicts predicted changes in signaling pathway activity based on the D10 vs D28 expression ratio changes. This phase is associated with osteoblast maturation, further increases in expression of secreted growth factors and matrix components, and the onset of matrix mineralization. The most notable changes during this interval are the activation of SMAD1/5/8 signaling downstream of BMP receptors and SMAD2/3 signaling from TGFβ receptors. Observed upregulation of the BMP receptors Bmpr1a (ALK3) and Bmpr2 (BMPR2) and the activin-like receptor Acvrl1 (ALK1), and SMAD2/3 likely contributes to the prediction of increased pathway activity. Upregulation of PTHrp/PTH1R and FGF2/FGF8 also suggests that GPCR and RTK signaling increase during this interval. Hence the data suggest that during osteoblastic differentiation of MC3T3-E1 cells, activation of β-catenin- and NFκB-mediated pathways occurs prior to the onset of TGFβ/BMP/SMAD-mediated signaling and a general activation of bone developmental signaling pathways.

**Fig 9.**
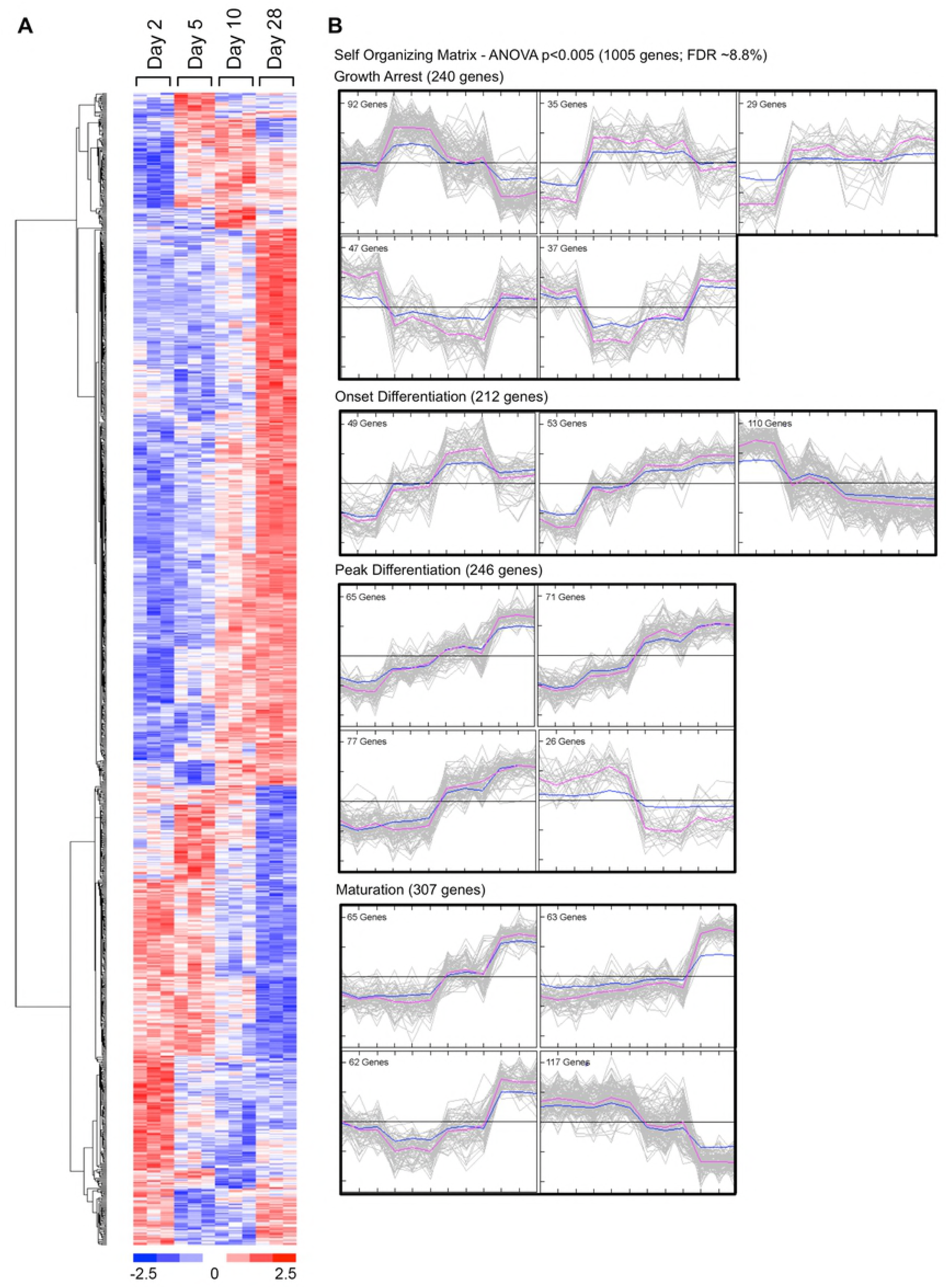
Changes in canonical signaling pathway activity in MC3T3-E1 cells between days 5 and 10. Expression ratios representing the changing abundance of 237 bone-related mRNA species in MC3T3-E1 cells between days 5 and 10 in culture were used to populate the IPA osteoarthritis pathway network and signaling pathway activation state was assessed using the IPA molecular pathway predictor tool. Observed upregulation and downregulation of mRNAs are shown in red and green, respectively, while predicted activation or inhibition of signaling intermediates and pathways are shown in orange and blue.

**Fig 10.**
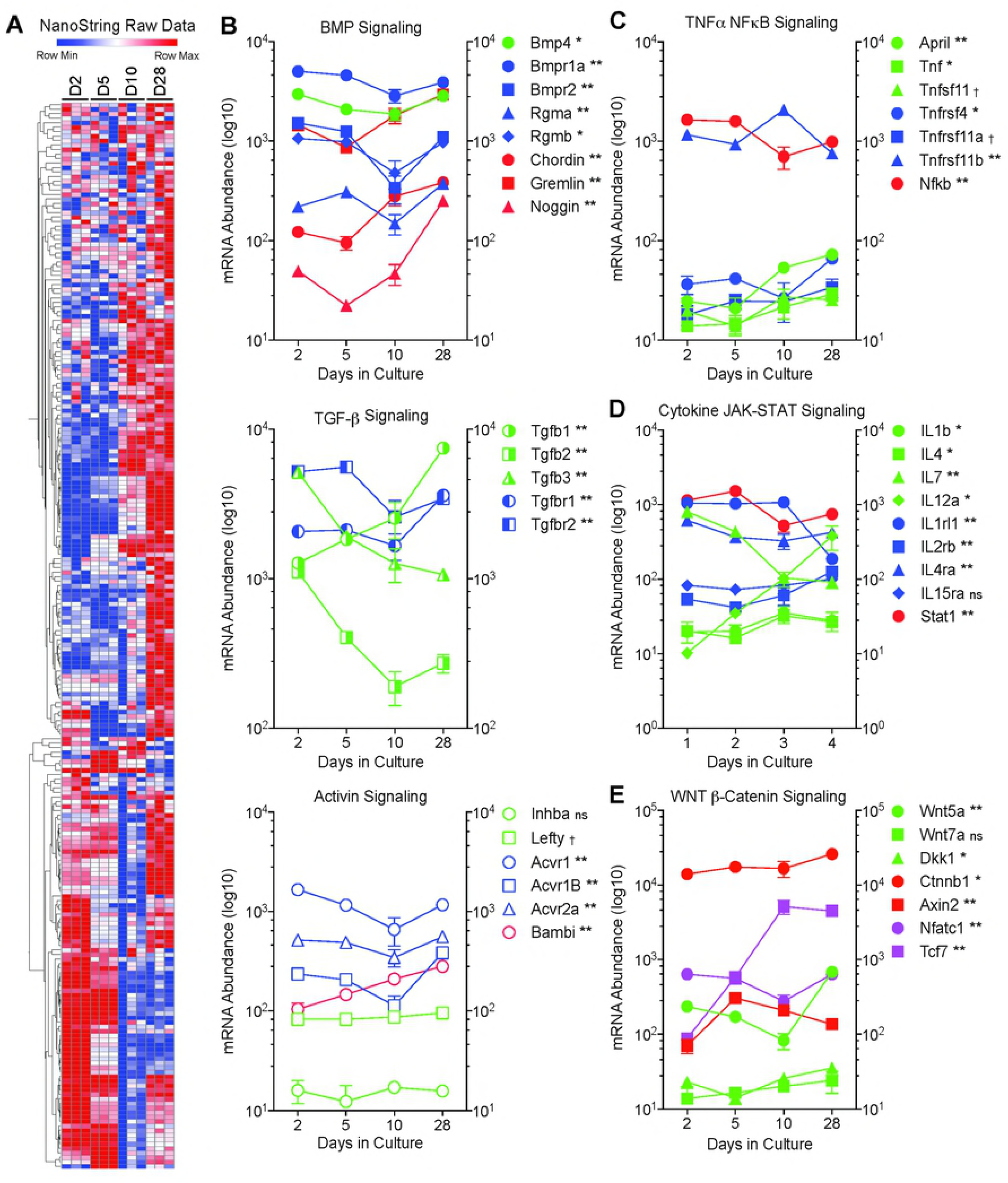
Changes in canonical signaling pathway activity in MC3T3-E1 cells between days 10 and 28. Expression ratios representing the changing abundance of 237 bone-related mRNA species in MC3T3-E1 cells between days 10 and 28 in culture were used to populate the IPA osteoarthritis pathway network and signaling pathway activation state was assessed using the IPA molecular pathway predictor tool. Observed upregulation and downregulation of mRNAs are shown in red and green, respectively, while predicted activation or inhibition of signaling intermediates and pathways are shown in orange and blue.

## Discussion

Complex biological processes like osteoblast development involve the coordinated regulation of multiple intracellular signaling pathways controlling gene expression. Thus, studies focusing on the contribution of any individual growth factor or pathway are invariably incomplete. Developing a more complete picture requires the use of “omics” approaches that capture as much information as possible about changes in intracellular signaling networks in as unbiased a manner as possible. Further complicating matters, bone remodeling *in vivo* is a continuous process wherein osteoblasts at all stages of development, from mesenchymal stem cell precursors to osteocytes are present, along with cells of the osteoclast lineage and other cell types [1,2]. As a result, studies performed on bone only provide a “snapshot” of the tissue average transcriptome that represents multiple cell types present in different proportions and differentiations states. In this study, we combined transcriptomics with bioinformatic geneset enrichment analysis to examine the temporal sequence of autocrine and paracrine signaling that regulates the differentiation of MC3T3-E1 cells, a well-characterized model of osteoblast development [7-10]. We employed two independently generated datasets, an “agnostic” DNA microarray dataset intended to provide a global overview of the evolving transcriptome and a “targeted” NanoString^™^ nCounter dataset focusing on genes involved in specific bone-related pathways. Our data complement other *in vitro* microarray studies of osteoblastic differentiation performed using different cell types, e.g. mesodermal progenitor cells, calvarial osteoblasts, osteocytes, periodontal ligament cells, and embryonic stem cells [21,26-30], or describing the effects of exogenous factors on osteoblast gene expression [31-34].

Although osteoblast differentiation *in vivo* is subject to regulation by numerous circulating factors [3], our results underscore the importance of cell autonomous autocrine/paracrine signaling. Key to the process is regulation of Runx2, the most upstream transcription factor in osteoblast differentiation [17,18], which regulates the expression of another critical transcription factor in bone, Sp7 [35]. Runx2 expression in osteoblasts is stimulated by an enhanceosome composed of Dlx5/6, Mef2, Tcf7, β-catenin, Sox5/6, Smad1, and Sp7, and in turn stimulates expression of bone matrix proteins including Spp1, Ibsp, and Bglap2, and autocrine factors including Ihh and Rankl [35,36]. Our data suggest that early MC3T3-E1 differentiation, between Days 2 and 5, is characterized by increasing expression of Runx2, Sp7, and β-catenin and upregulation of IHH, TNFα, and IL1β at a time where TGFβ/BMP/SMAD signaling is still relatively suppressed despite increasing expression of TGFβ and TGFβ receptors. WNT signaling, which cooperates with TGFβ in a positive regulatory loop by inducing Runx2-dependent transcription of TGFβ1 receptor [37,38], appeared most active early and to wane as TGFβ/BMP pathway activity increased, consistent with a role for WNT signaling in the induction of TGFβ signaling. The central role of the TGFβ/BMP axis in regulating mesenchymal stem cell differentiation into cartilage and bone is well established, as both canonical SMAD-dependent and non-canonical p38MAPK signaling downstream of these receptors converge on Runx2 to promote differentiation [19-22]. Moreover, in bone TGFβ/BMP in extensive cross talk with other signaling pathways [4,23]. Of these, the activity of several, including Hedgehog, FGF2, interleukins, TNFα/RANKL and interferon-γ, appeared to increase in parallel with TGFβ/BMP during MC3T3-E1 cell differentiation. Hedgehog signaling, acting through Gli family transcription factors, promotes the expression of BMP2, and IHH has been shown to be required for osteogenesis *in vitro* [39,40]. FGF2 regulates expression of PC1, the primary enzymatic generator of pyrophosphate in mineralizing cells, by direct regulation of Runx2, suggesting that TGFβ/BMP and FGF2 signaling cooperate to promote matrix mineralization later in differentiation [41]. TNFα plays many roles in bone, inhibiting osteoblast differentiation and collagen synthesis (42,43), promoting osteoblast apoptosis (44), while directly stimulating osteoclast formation independent of RANKL signaling through an IL1-dependent mechanism (45,46). Conversely, interferon-γ opposes IL1 and TNFα mediated bone resorption, but produces additive inhibition of bone collagen synthesis (47). Thus our network analysis, demonstrating simultaneous changes in the TGFβ/BMP pathways that favoring osteoblast differentiation and survival, the TNFα pathway that inhibits differentiation and favors apoptosis, and the interferon-γ pathway that inhibits ongoing collagen synthesis, illustrates the complexity of osteoblast development and maturation that occur in the setting of opposing autocrine signaling loops.

While gene array technology is a powerful tool for determining the transcriptional basis of changing developmental or pharmacological processes, the resulting datasets are both too complex and too error prone to reliably base conclusions on casual inspection [5,6]. Metabolic pathways analysis overcomes several of these limitations. By basing conclusions on the number and magnitude of expression changes across gene clusters, rather than individual genes, it decreases the probability of false discovery, while simultaneously providing a quantitative measure of the probability of change in a given signaling network. Our analyses illustrate several of the advantages and disadvantages of this approach. The murine Operon V2.0 cDNA arrays employed in this study did not provide genome-wide coverage of changes in mRNA abundance. The NanoString^™^ nCounter system provides information only about rationally chosen transcripts. Hence, some information is missing. Moreover, transcriptomic datasets in general are limited in their ability to discriminate changes in cellular metabolism simply because important pathway components may not be regulated at the transcriptional level, rendering them “invisible” in gene array experiments. Bioinformatic tools, such as IPA, that “infer” changes in upstream pathway activity based on observed changes in network components, provide a means to translate incomplete transcriptomic datasets into a more complete picture of metabolic activity. In our study, we took two independently generated sets of mRNA samples from differentiating MC3T3-E1 cells and employed two different approaches to pathways analysis. Close inspection of the data (S1 and S2 Tables) shows that the relatively stringent statistical filter applied to the microarray dataset to define significant change failed to capture factors that were seen in the NanoString^™^ assay, and conversely, that the targeted NanoString^™^ Code Set missed significant changes in factors that were detected with the broader coverage provided by the microarrays. It is also noteworthy that some of the interval expression ratios of individual factors were seen to change in opposite directions in the two datasets, such that focusing on the abundance of individual factors might lead to different conclusions. Yet the remarkable degree of similarity in the pathways analysis, which weighs changes across entire networks to predict pathway activation, suggests the approach is both reliable and robust enough to tolerate a substantial amount of “noise” in the raw data. Thus, starting from incomplete datasets, we were able to extract temporal changes in the autocrine/paracrine signaling networks that influence osteoblast differentiation *in vitro*, and find evidence of pathway cross talk at the transcriptional level.

## Acknowledgements

Supported by NIH/NIGMS (R01 GM095497 and R35 GM126955) and the United States Depar™ent of Veterans Affairs (I01 BX003188). Some bioinformatic analyses were conducted through the Medical University of South Carolina (MUSC)

Proteogenomics Facility that was supported by NIH/NIGMS (P30 GM103342 and P20 GM103499) and MUSC’s Office of the Vice President for Research. The contents of this article do not represent the views of the Depar™ent of Veterans Affairs or the United States Government.

## Author Contributions

Conceptualization: DGP, LML. Formal Analysis: LML, JLB. Funding Acquisition: LML. Investigation: MSD, HME-S, DGP, CJH, KMR. Writing: JLB, HME-S, DGP, CJH, LML.

## Supporting Information Captions

**S1 Fig. Comparison of WNT/β-catenin network activity between Days 2 and 5 of MC3T3-E1 cell differentiation predicted from the microarray and NanoString^™^ datasets.** Observed Day 2 to Day 5 changes in expression ratios were used to predict WNT/β-catenin pathway activity using the IPA molecular activity predictor tool. **A.** Pathway activity prediction based on the microarray dataset. **B.** Pathway activity based on the NanoString^™^ dataset. Observed increases (*red*) and decreases (*green*) in mRNA abundance are indicated, as are predicted activation (*orange*) and inhibition (*blue*) of downstream targets.

**S2 Fig. Comparison of TGFβ/BMP network activity between Days 10 and 28 of MC3T3-E1 cell differentiation predicted from the microarray and NanoString^™^ datasets.** Observed Day 10 to Day 28 changes in expression ratios were used to predict TGFβ/BMP pathway activity using the IPA molecular activity predictor tool. **A.** Pathway activity prediction based on the microarray dataset. **B.** Pathway activity based on the NanoString^™^ dataset. Observed increases (*red*) and decreases (*green*) in mRNA abundance are indicated, as are predicted activation (*orange*) and inhibition (*blue*) of downstream targets.

**S3 Fig. Comparison of NFκB network activity between Days 10 and 28 of MC3T3-E1 cell differentiation predicted from the microarray and NanoString^™^ datasets.** Observed Day 10 to Day 28 changes in expression ratios were used to predict NFκB pathway activity using the IPA molecular activity predictor tool. **A.** Pathway activity prediction based on the microarray dataset. **B.** Pathway activity based on the NanoString^™^ dataset. Observed increases (*red*) and decreases (*green*) in mRNA abundance are indicated, as are predicted activation (*orange*) and inhibition (*blue*) of downstream targets.

**S4 Fig. Comparison of STAT3 network activity between Days 10 and 28 of MC3T3-E1 cell differentiation predicted from the microarray and NanoString^™^ datasets.** Observed Day 10 to Day 28 changes in expression ratios were used to predict STAT3 pathway activity using the IPA molecular activity predictor tool. **A.** Pathway activity prediction based on the microarray dataset. **B.** Pathway activity based on the NanoString^™^ dataset. Observed increases (*red*) and decreases (*green*) in mRNA abundance are indicated, as are predicted activation (*orange*) and inhibition (*blue*) of downstream targets.

**S1 Table. NanoString^TM^ nCounter expression data for 237 bone-related transcripts.** Gene symbol, accession number, annotation, NanoString^™^ probe ID, and mRNA abundance data are shown for triplicate determinations at each of four time points.

**S2 Table. Operon V2.0 microarray expression data for 1005 significantly-regulated transcripts.** Gene symbol, accession number, gene name, mRNA abundance data, and z-standardized expression values are shown for triplicate determinations at each of four time points.

**S3 Table. IPA Disease or Function analysis of significantly-regulated transcripts identified by microarray.** Disease or function annotation, -log(p value), activation z-score, number and name of pathway molecules are shown for all functions with activation z-score >1 or <-1 in the D2 vs D5, D5 vs D10, and D10 vs D28 pairwise comparisons.

**S4 Table. IPA Upstream Regulator analysis of significantly-regulated transcripts identified by microarray.** Gene symbol and activation z-score are shown for all upstream regulators with activation z-score >2 or <-2 in the D2 vs D5, D5 vs D10, and D10 vs D28 pairwise comparisons.

**S5 Table. IPA Canonical Pathways analysis of significantly-regulated transcripts identified by microarray.** Canonical Pathway name, -log(pvalue), activation z-score, and observed pathway molecules are shown for predicted regulated pathways in the D2 vs D5, D5 vs D10, and D10 vs D28 pairwise comparisons.

**S6 Table. IPA Upstream Regulator analysis of the NanoString dataset of bone-related genes.** Gene symbol and activation z-score are shown for all upstream regulators with activation z-score >2 or <-2 in the D2 vs D5, D5 vs D10, and D10 vs D28 pairwise comparisons.

**S7 Table. IPA Canonical Pathways analysis of the NanoString dataset of bone-related genes.** Canonical Pathway name, -log(pvalue), activation z-score, and observed pathway molecules are shown for predicted regulated pathways in the D2 vs D5, D5 vs D10, and D10 vs D28 pairwise comparisons.

